# System-level analyses of keystone genes required for mammalian tooth development

**DOI:** 10.1101/869065

**Authors:** Outi Hallikas, Rishi Das Roy, Mona M. Christensen, Elodie Renvoisé, Ana-Marija Sulic, Jukka Jernvall

## Abstract

When a null mutation of a gene causes a complete developmental arrest, the gene is typically considered essential for life. Yet, in most cases null mutations have more subtle effects on the phenotype. Here we used the phenotypic severity of mutations as a tool to examine system-level dynamics of gene expression. We classify genes required for the normal development of the mouse molar into different categories that range from essential to subtle modification of the phenotype. Collectively, we call these the developmental keystone genes. Transcriptome profiling using microarray and RNAseq analyses of patterning stage mouse molars show highly elevated expression levels for genes essential for the progression of tooth development, a result reminiscent of essential genes in single cell organisms. Elevated expression levels of progression genes were also detected in developing rat molars, suggesting evolutionary conservation of this system-level dynamics. Single-cell RNAseq analyses of developing mouse molars reveal that even though the size of the expression domain, measured in number of cells, is the main driver of organ-level expression, progression genes show high cell-level transcript abundances. Progression genes are also upregulated within their pathways, which themselves are highly expressed. In contrast, a high proportion of the genes required for normal tooth patterning are secreted ligands that are expressed in fewer cells than their receptors and intracellular components. Overall, even though expression patterns of individual genes can be highly different, conserved system-level principles of gene expression can be detected using phenotypically defined gene categories.

## 1| INTRODUCTION

Much of the functional evidence for the roles of developmental genes comes from natural mutants or experiments in which the activity of a gene is altered. Most often these experiments involve deactivation, or a null mutation where the production of a specific gene product is prevented altogether. In the cases where development of an organism is arrested, the specific gene is considered to be absolutely required or essential for development (Amsterdam et al., 2004; Dickinson et al., 2016). Through a large number of experiments in different organisms, an increasingly nuanced view of developmental regulation has emerged showing that some genes appear to be absolutely required, whereas others may cause milder effects on the phenotype (Brown et al., 2018; Bogue et al., 2018). Yet, there are a large number of genes that, despite being dynamically regulated during individual organ development, have no detectable phenotypic effect when null mutated.

Within the framework of distinct phenotypic outcomes of gene deactivation it can be argued that there is a gradation from developmentally ‘more essential’ to ‘less essential’ genes. Collectively, these can be considered to be analogous to the keystone species concept used in ecology (Paine, 1969; Terborgh, 1986). These genes, which can be called ‘developmental keystone genes’, are not necessarily essential for development. Rather, compared to all the genes, developmental keystone genes exert a disproportional effect on the phenotype.

As large-scale analyses of transcriptomes produce expression profiles for individual organs at the organ and single-cell level, it is now possible to address whether there might be any system-level differences between the regulation of essential and other keystone genes during organogenesis. Here we address such differences using the mammalian tooth. Especially the development of the mouse molar is well characterized, with over 70 genes that are known to be individually required for normal tooth development (Bei, 2009; Nieminen, 2009; Harjunmaa et al., 2012). The dynamic expression patterns and detailed effects of null mutations of these genes are also exceedingly well characterized, ranging from a complete developmental arrest to relatively mild modifications of morphology, or defects in the mineralized hard tissue (Nieminen, Pekkanen, Aberg, & Thesleff, 1998; Nieminen, 2009; Harjunmaa et al., 2012). By classifying these genes into different categories (Figure 1), we investigated whether there are any differences in the expression of different gene categories.

**Figure 1.**
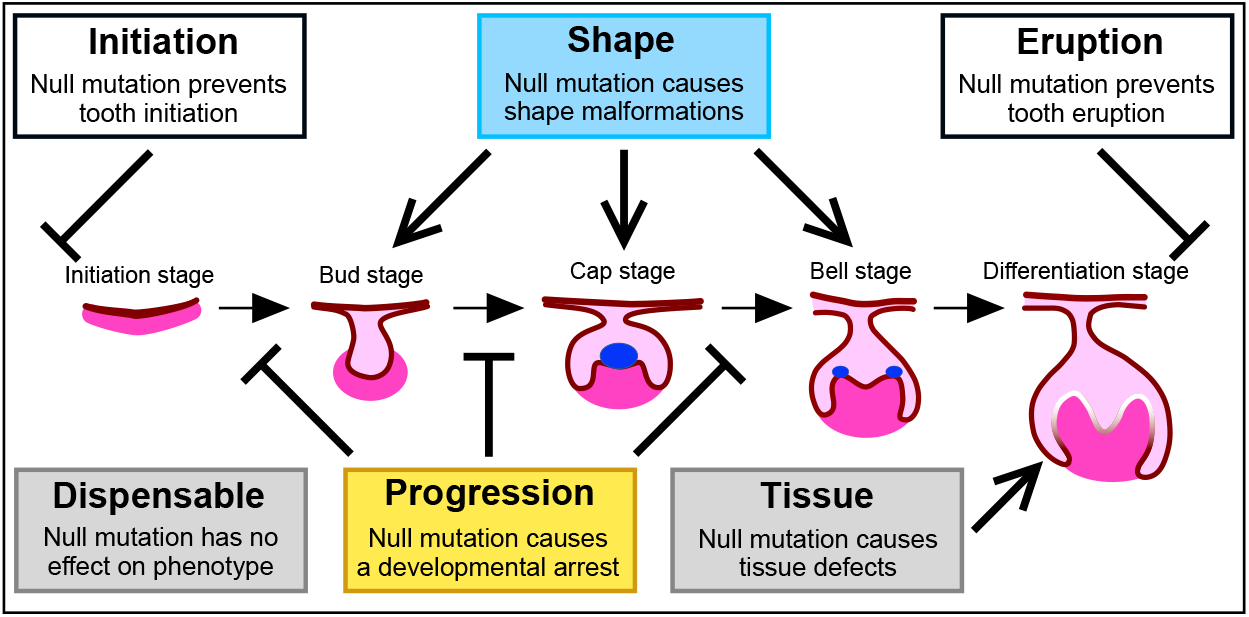
Keystone gene categories of tooth development. Mouse molar development progresses from initiation and patterning to formation of the hard tissues and eruption. These steps are mediated by reciprocal signaling between epithelium (pink) and mesenchyme (magenta). A central step in the patterning is the formation of the epithelial signaling center, the primary enamel knot (blue oval inside the cap stage tooth). Several genes are known to be required for the developmental progression and regulation of the shape around the time of cap stage, and here we focused mainly on transcriptomes in the bud, and cap stage molars. Expression of progression and shape category genes were compared to tissue and dispensable genes, as also to other developmental process genes. Fewer initiation and eruption category genes are known, and they were excluded from the analyses. For listing of the genes, see Appendix S1 and Table S1.

## 2| MATERIALS AND METHODS

### 2.1| Classification of developmental keystone genes

Here our main focus is on a critical step in tooth development, namely the formation of the cap stage tooth germ (Figure 1). At this stage the patterning of tooth crown begins, and the effects of experimental modifications in several signaling pathways first manifest themselves around this time of development (Jernvall & Thesleff, 2012). To classify the studied genes, we divide them into different categories based on our analysis of published experiments. Our classification scheme is based on the phenotype of the mouse where that gene is knocked out. Thus, the classification of each gene is based on in vivo experiments. Operationally, our classification applies only to our organ of interest even though classification following the same logic could be done for any organ. This single organ focus also means that genes that have no effect in one organ may be critical for the development of another organ. Because many developmental genes function in multiple organs and stages during development, full mutants of several genes are lethal before tooth development even begins. Therefore, when available, we also used information on the tooth phenotypes of conditional mutant mice. We note that whereas the keystone gene terminology has also been considered within the context of their effects on ecosystems (Skovmand et al., 2018), here we limit the explanatory level to a specific organ system.

The first category is the **progression category** containing essential genes that cause a developmental arrest of the tooth when null mutated (Figure 1, genes with references in Appendix S1). The second set of genes belongs to the **shape category** and they alter the morphology of the tooth when null mutated. Unlike the null-mutations of progression genes, many shape gene mutations cause subtle modifications of teeth that remain functional, hence these genes are not strictly essential for tooth development. The third category is the **tissue category** and null mutations in these genes cause defects in the tooth hard tissues, enamel and dentine. Both the progression and shape categories include genes that are required for normal cap-stage formation. In contrast, the tissue category is principally related to the formation of extracellular matrix and these genes are known to be needed much later in the development (Nieminen at al. 1998). Because there is more than a five-day delay from the cap-stage to matrix secretion in the mouse molar, here we considered the tissue category as a control for the first two categories.

Additionally, we compiled a second control set of developmental genes that, while expressed during tooth development, are reported to lack phenotypic effects when null mutated (Table S1). This **dispensable category** is defined purely within our operational framework of identifiable phenotypic effects and we do not imply that these genes are necessarily unimportant even within the context of tooth development. Many genes function in concert and the effects of their deletion only manifest when mutated in combinations (also known as synthetic mutations). Some dispensable genes may function in combinations even though no such evidence exists as yet. We identified five such redundant pairs of paralogous genes and a single gene whose null phenotype surfaces in heterozygous background of its paralogue. Altogether these 11 genes were tabulated separately as a **double category**. In the progression, shape, tissue, and dispensable categories we tabulated 15, 28, 27, and 100 genes respectively (Figure 1, Appendix S1, Table S1). While still limited, these genes should represent a robust classification of validated experimental effects. We note that these groupings do not exclude the possibility that a progression gene, for example, can also be required for normal hard tissue formation. Therefore, the keystone gene categories can be considered to reflect the temporal order in which they are first required during odontogenesis. Moreover, many of the 100 dispensable category genes could belong to the double category, but finding them would require testing close to 5 000 transgenic combinations to test.

In addition to the categories studied here, there are genes required for the initiation of tooth development, of which many are also potentially involved in tooth renewal. Because the phenotypic effect of these initiation genes on tooth development precedes the visible morphogenesis, and the phenotype might include complete lack of cells of the odontogenic lineage, we excluded these genes from our analyses. Similarly, we excluded genes preventing tooth eruption with no specific effect on the tooth itself (Appendix S1).

To examine our gene categories in the context of whole transcriptomes, we compared the expression levels with all the developmental-process genes (GO:0032502; Ashburner et al., 2000), as also with all the other protein coding genes. The developmental-process genes with GO term “GO:0032502” and experimental evidence codes were obtained from R package “org.Mm.eg.db” (Carlson, 2019). Only curated RefSeq genes are used in this study. All tabulations are found in Appendix S1 and Table S1, and S4. For the analyses of rat teeth, the classification of mouse genes was transferred to one-to-one orthologs of rat genome. The data of orthologs were downloaded from Ensembl server using R package “biomart” (Durinck et al., 2005).

For the analyses of pathways, we created a manually curated list of genes in the six key pathways (Wnt, Tgfβ, Fgf, Hh, Eda, Notch) and allocated the genes into these pathways where appropriate. Genes were also classified as ‘ligand’ (signal), ‘receptor’, ‘intracellular molecule’, ‘transcription factor’ or ‘other’. Because these kinds of classifications are not always trivial as some biological molecules have multiple functions in the cell, we used the inferred primary role in teeth.

### 2.2| Ethics statement

All mouse and rat studies were approved and carried out in accordance with the guidelines of the Finnish national animal experimentation board under licenses KEK16-021, ESAVI/2984/04.10.07/2014 and ESAV/2363/04.10.07/2017.

### 2.3| Dissection of teeth

Wild type tooth germs were dissected from mouse embryonic stages corresponding to E13, E14 and E16 molars. For bulk and single-cell RNAseq we used C57BL/6JOlaHsd mice, and for microarray we used NMRI mice. The wild type rat tooth germs were dissected from DA/HanRj rat embryonic stages E15 and E17, which correspond morphologically to E13 and E14 mouse molars (mouse and rat molars are relatively similar in shape). Minimal amount of surrounding tissue was left around the tooth germ, at the same time making sure that the tooth was not damaged in the process. The tissue was immediately stored in RNAlater (Qiagen GmbH, Hilden, Germany) in −80°C for RNAseq or in TRI Reagent (Merck, Darmstadt, Germany) in - 80°C for microarray. For microarray, a few tooth germs were pooled for each sample and five biological replicas were made. For RNAseq, each tooth was handled individually. Seven biological replicates were made for mouse and five biological replicates for rat. Numbers of left and right teeth were balanced.

### 2.4| RNA extraction

The tooth germ was homogenised into TRI Reagent (Merck, Darmstadt, Germany) using Precellys 24-homogenizer (Bertin Instruments, Montigny-le-Bretonneux, France). The RNA was extracted by guanidium thiocyanate-phenol-chloroform method and then further purified by RNeasy Plus micro kit (Qiagen GmbH, Hilden, Germany) according to manufacturer’s instructions. The RNA quality was assessed for some samples with 2100 Bioanalyzer (Agilent, Santa Clara, CA) and all the RIN values were above 9. The purity of RNA was analysed by Nanodrop microvolume spectrophotometer (ThermoFisher Scientific, Waltham, USA). RNA concentration was measured by Qubit 3.0 Fluorometer (ThermoFisher Scientific, Waltham, USA). The cDNA libraries were prepared with Ovation Mouse RNAseq System or Ovation Rat RNAseq System (Tecan, Zürich, Schwitzerland).

### 2.5| Bulk RNA expression analysis

Gene expression levels were measured both in microarray (Affymetrix Mouse Exon Array 1.0, GPL6096) and RNAseq (platforms GPL19057, Illumina NextSeq 500). The microarray gene signals were normalized with aroma.affymetrix (Bengtsson, Irizarry, Carvalho, & Speed, 2008) package using Brainarray custom CDF (Version 23, released on Aug 12, 2019) (Dai et al., 2005). The RNAseq reads (84 bp) of mouse and rat were evaluated and bad reads are filtered out using FastQC (Andrews et al., 2012), AfterQC (Chen et al., 2017) and Trimmomatic (Bolger, Lohse, & Usadel, 2014). This resulted on average 63 million reads per mouse sample and 45 million reads per rat sample. Good reads for mouse and rat were aligned with STAR (Dobin et al., 2013) to GRCm38 (mm10/Ensembl release 90) and Rnor_6.0 (Ensembl release 101), respectively. Counts for each gene were performed by HTSeq (Anders, Pyl, & Huber, 2015) tool. Results are shown without normalization of gene expression based on gene length as it does not change the pattern of results. On average 85% of reads were uniquely mapped to the genome.

### 2.6| Single-cell RNA sequencing

Single cell RNA sequencing was performed on mouse E14 cap stage tooth cells. The teeth were dissected as described above. Each tooth was processed individually in the single-cell dissociation. In total 4 teeth were analyzed. Each tooth germ was treated with 0.1 mg/ml liberase (Roche, Basel, Schwitzerland) in Dulbecco’s solution for 15 min at 28°C in shaking at 300 rpm followed by gentle pipetting to detach the mesenchymal cells. Then the tissue preparation was treated with TrypLE Select (Life Technologies, Waltham, USA) for 15 min at 28°C in shaking at 300 rpm followed by gentle pipetting to detach the epithelial cells. The cells were washed once in PBS with 0,04% BSA. The cells were resuspended in 50 μl PBS with 0.04% BSA. We used the Chromium single cell 3’ library & gel bead Kit v3 (10x Genomics, Pleasanton, USA). In short, all samples and reagents were prepared and loaded into the chip. Then, Chromium controller was used for droplet generation. Reverse transcription was conducted in the droplets. cDNA was recovered through emulsification and bead purification. Pre-amplified cDNA was further subjected to library preparation. Libraries were sequenced on an Illumina Novaseq 6000 (Illumina, San Diego, USA). All the sequencing data are available in GEO under the accession number GSE142201.

### 2.7| Data analysis

For scRNAseq, 10x Genomics Cell Ranger v3.0.1 pipelines were used for data processing and analysis. The “cellranger mkfastq” was used to produce fastq files and “cellranger count” to perform alignment, filtering and UMI counting. Alignment was done against mouse genome GRCm38/mm10. The resultant individual count data were finally aggregated with “cellranger aggr”. Further, the filtered aggregated feature-barcode matrix was checked for quality and normalization using R package Seurat (Stuart et al., 2019). Only cells with ≥ 20 genes and genes expressed in at least 3 cells were considered for all the downstream analysis. For a robust set of cells for the expression level calculations, we limited the analyses to 30930 cells that had transcripts from 3000 to 9000 genes (7000 to 180000 unique molecular identifiers) with less than 10% of the transcripts being mitochondrial. For comparison with bulk RNAseq data (Figs 3), single-cell data was normalized with DeSeq2 (Love, Huber, & Anders, 2014) together with the corresponding bulk RNAseq samples, and median expression levels were plotted. The average cell-level expression (Figure 4B) of a gene *X* was calculated as

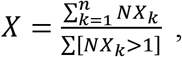

where *NX_k_* is normalized expression of gene *X* in cell *k* and the denominator is the count of cells with non-zero reads. All statistical tests corresponding to Tables S3 and S4 were performed using R package “rcompanion” (Mangiafico, 2019) and custom R scripts.

## 3| RESULTS

### 3.1| Especially progression genes show elevated expression at the onset of tooth patterning

For a robust readout of gene expression profiles, we first obtained gene expression levels using both microarray and RNAseq techniques from E13 (bud stage) and E14 (cap stage) mouse molars (Materials and Methods). From dissected tooth germs we obtained five microarray and seven RNAseq replicates for both developmental stages. The results show that especially the progression category genes (genes required for the progression of tooth development) are highly expressed during E13 compared to the control gene sets (tissue, dispensable, and developmental-process categories, *p* values range from 0.0003 to 0.0426 for RNAseq and microarray experiments, tested using random resampling, for details and all the tests, see Materials and Methods, Figure 2, Tables S2, S3). Comparable differences are observed in E14 molars (*p* values range from 0.0000 to 0.0466, Figure 2, Tables S2, S3).

**Figure 2.**
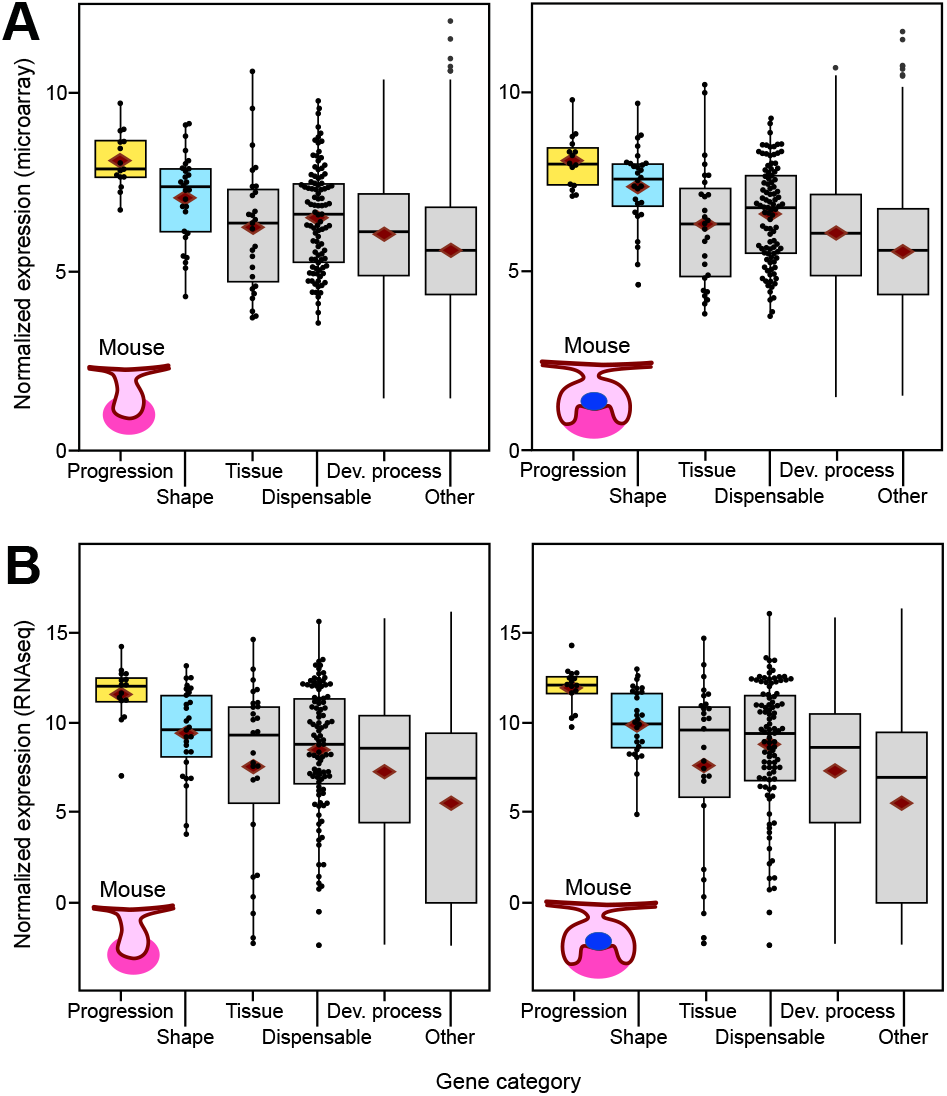
Bud and cap stage mouse molars show elevated expression of progression and shape genes. (A) Microarray and (B) RNAseq analyses of the transcriptomes of E13 and E14 molars show highly elevated expression of progression category and moderately elevated shape category genes (for tests, see Table S3). Tissue category genes are involved in dentine and enamel formation that begin at birth, around six days from the E14 cap stage. The number of genes having RNAseq expression data in each category are 15, 28, 27, 100, 4106, and 16165 for progression, shape, tissue, dispensable, developmental process, and other, respectively. The corresponding numbers for the microarray data are 15, 28, 27, 98, 3983, and 14825. Boxes enclose 50% of observations; the median and mean are indicated with a horizontal bar and diamond, respectively, and whiskers extend to last values within 1.5 interquartiles. Individual data points are shown for the smaller categories.

In general, the expression differences between progression and tissue categories appear greater than between progression and dispensable categories (*p* values range from 0.0028 to 0.0379 and 0.0059 to 0.0466, respectively, Table S3), suggesting that some of the genes in the dispensable category may still play a functional role in tooth development. In our data we have 11 genes that cause a developmental arrest of the tooth when double mutated (Appendix S1). The expression level of this double-mutant category shows incipient upregulation compared to that of the developmental-process category (*p* values range from 0.0322 to 0.1637 Table S3), but not when compared to the tissue or dispensable categories (*p* values range from 0.0978 to 0.5010, Table S3). Therefore, it is plausible, based on the comparable expression levels between double and some of the dispensable category genes, that several of the genes in the dispensable category may cause phenotypic effects when mutated in pairs.

Even though expression levels of the shape category genes (genes required for normal shape development) are lower than that of the progression category (Figure 2), at least the E14 microarray data suggests elevated expression levels relative to all the other control categories (*p* values range from 0.0001 to 0.0901, Table S3). The moderately elevated levels of expression by the shape category genes could indicate that they are required slightly later in development, or that the most robust upregulation happens only for genes that are essential for the progression of the development. The latter option seems to be supported by a RNAseq analysis of E16 molar, showing only slight upregulation of shape category genes in the bell stage molars (Table S3).

### 3.2| Transcriptomes of developing rat molars show elevated expression of the progression genes

Because our gene categories were based on experimental evidence from the mouse, we also tested whether comparable expression levels can be detected for the same genes in the rat. Evolutionary divergence of *Mus-Rattus* dates back to the Middle Miocene (Kimura et al., 2015), allowing a modest approximation of conservation in the expression levels. Examination of bud (E15) and cap (E17) stage RNAseq of rat molars shows comparable upregulation of progression and shape category genes as in the mouse (Figure 3, Table S2, S3). Considering also that many of the null mutations in keystone gene in the mouse are known to have comparable phenotypic effects in humans (Nieminen, 2009), our keystone gene categories and analyses are likely to apply to mammalian teeth in general.

**Figure 3.**
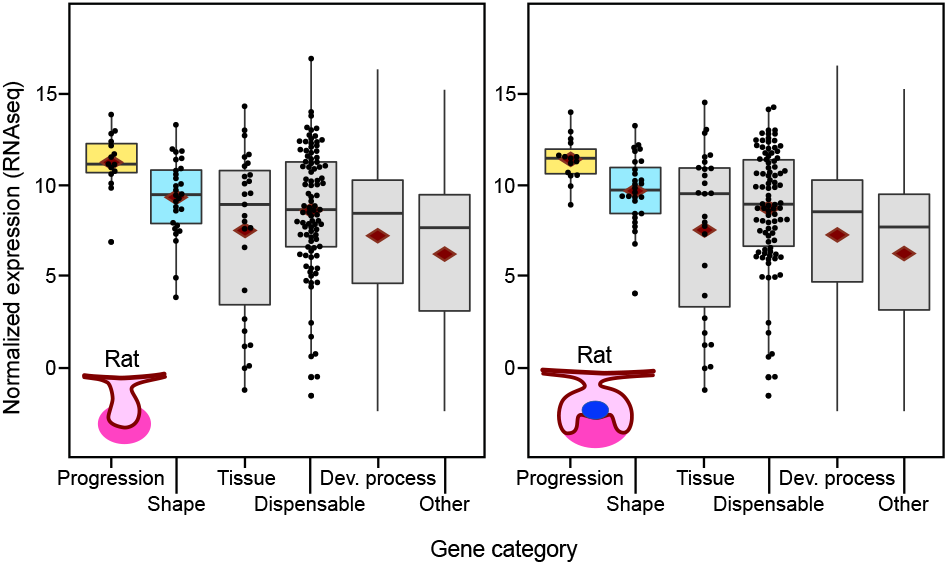
Bud and cap stage rat molars show elevated expression of keystone genes. RNAseq analyses of the transcriptomes of E15 and E17 rat molars show highly elevated expression of progression category and moderately elevated shape category genes (for tests, see Table S3). The numbers of genes having RNAseq expression data in each category are 15, 28, 27, 95, 3843, and 12473 for progression, shape, tissue, dispensable, developmental process, and other, respectively. Boxes enclose 50% of observations; the median and mean are indicated with a horizontal bar and diamond, respectively, and whiskers extend to last values within 1.5 interquartiles. Individual data points are shown for the smaller categories.

One complication of our expression level analyses is that these have been done at the whole organ level. Because many of the genes regulating tooth development are known to have spatially complex expression patterns within the tooth (Nieminen at al. 1998), cell-level examinations are required to decompose the patterns within the tissue.

### 3.3| Single-cell RNAseq reveals cell-level patterns of keystone genes

Tooth development is punctuated by iteratively forming epithelial signaling centers, the enamel knots. The first, primary enamel knot, is active in E14 mouse molar and at this stage many genes are known to have complex expression patterns. Some progression category genes have been reported to be expressed in the enamel knot, whereas others have mesenchymal or complex combinatorial expression patterns (Jernvall & Thesleff, 2012; Nieminen et al., 1998). To quantify these expression levels at the cell-level, we performed a single-cell RNAseq (scRNAseq) on E14 mouse molars (Materials and Methods). We focused on capturing a representative sample of cells by dissociating each tooth germ without cell sorting (*n* = 4). After data filtering, 7000 to 8811 cells per tooth were retained for the analyses, providing 30930 aggregated cells for a relatively good proxy of the E14 mouse molar (Materials and Methods).

First we examined whether the scRNAseq produces comparable expression levels to our previous analyses. For the comparisons, the gene count values from the cells were summed up and treated as bulk RNAseq data (Figure 4A, Materials and Methods). We analyzed the expression levels of different gene categories as in the mouse bulk data (Figure 2) and the results show a general agreement between the experiments (Figure 2, 4B). As in the previous analyses (Table S3), the progression category shows the highest expression levels compared to the control gene sets (*p* values range from 0.0071 to 0.0310, Table S3). Although the mean expression of the shape category is intermediate between progression and control gene sets, scRNAseq shape category is not significantly upregulated in the randomization tests (*p* values range from 0.7788 to 0.9968). This pattern reflects the bulk RNAseq analyses (for both mouse and rat) while the microarray analysis showed slightly stronger upregulation, suggesting subtle differences between the methodologies (the used mouse strain was also different in the microarray experiment).

**Figure 4.**
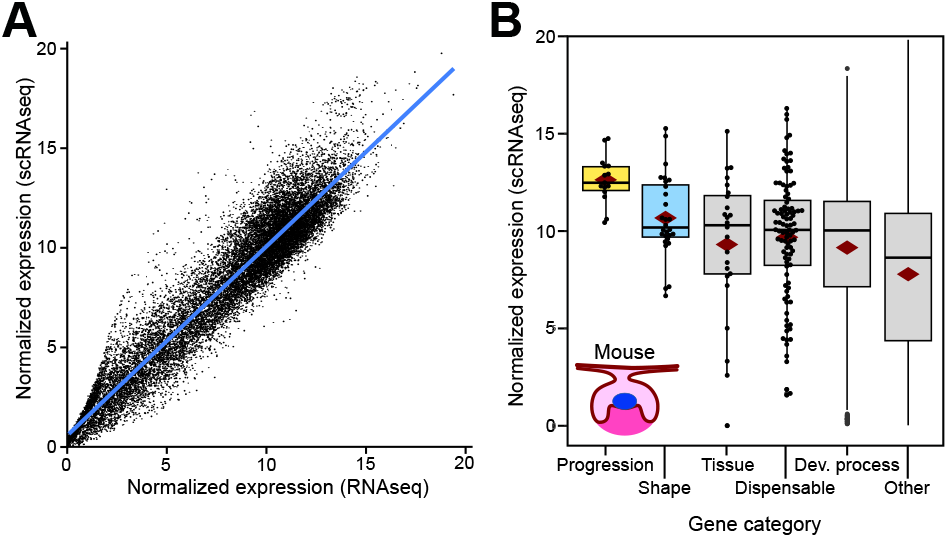
Single-cell RNAseq data reflect the bulkRNA analyses. (A) Bulk and scRNAseq expression levels show overall correspondence in tissue level expression in E14 mouse molar. (B) Progression category genes show the strongest upregulation whereas shape category is intermediate between the progression and other categories (for tests, see Table S3). The number of genes having expression data in each category are 15, 28, 25, 99, 3771, and 16362 for progression, shape, tissue, dispensable, developmental process, and other, respectively. Boxes in (B) enclose 50% of observations; the median and mean are indicated with a horizontal bar and diamond, respectively, and whiskers extend to last values within 1.5 interquartiles. Individual data points are shown for the smaller categories.

Unlike the bulk transcriptome data, the scRNAseq data can be used to quantify the effect of expression domain size on the overall expression level of a gene. The importance of expression domain size is well evident in the scRNAseq data when we calculated the number of cells that express each gene (Materials and Methods). The data shows that the overall tissue level gene expression is highly correlated with the cell population size (Figure 5A). In other words, the size of the expression domain is the key driver of expression levels measured at the whole tissue level.

**Figure 5.**
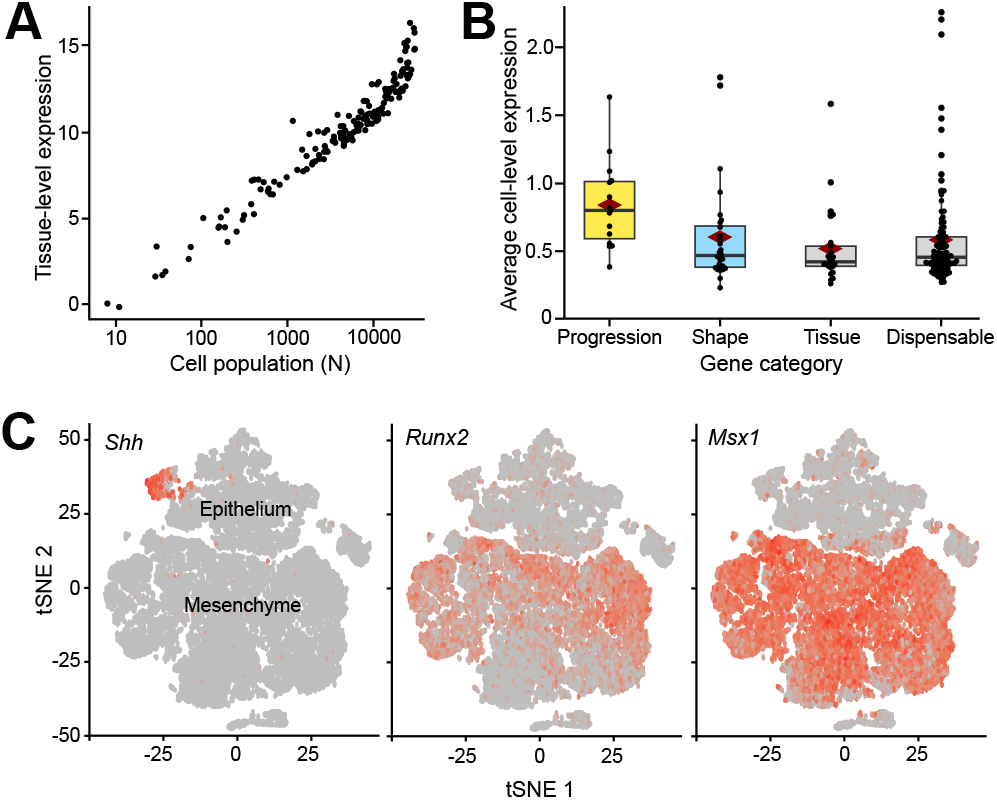
Expression domain size and high cell-level transcript abundance of progression genes. (A) The size of the expression domain, measured as the number of cells, is the key driver of organ level expression (scRNAseq on E14 mouse molar, plotted for progression, shape, tissue, and dispensable category genes). (B) Progression category shows high cell-level expression or transcript abundance, indicating high expression relative to the expression domain size. The *p* values for progression, shape, and tissue categories compared to dispensable category are 0.0008, 0.7009, 0.3413, respectively (one-tailed significance levels obtained using 10 000 permutations). (C) tSNE plots showing diverse expression patterns of three different progression category genes in the scRNAseq data. Clusters containing epithelial and mesenchymal cells are marked (note the limited presence of mesenchymal marker *Msx1* transcripts in the epithelium, agreeing with previous reports by Coudert et al., 2005). Boxes in (B) enclose 50% of observations; the median and mean are indicated with a horizontal bar and diamond, respectively, and whiskers extend to last values within 1.5 interquartiles.

To examine the cell level patterns further, we calculated the mean transcript abundances for each gene for the cells that express that gene (see Material and Methods). This metric approximates the cell-level upregulation of a particular gene, and is thus independent of the size of the expression domain. We calculated the transcript abundance values for progression, shape, tissue, double, and dispensable category genes in each cell that expresses any of those genes. The resulting mean transcript abundances were contrasted to that of the dispensable category (Materials and Methods). The results show that the average transcript abundance is high in the progression category whereas the other categories show roughly comparable transcript abundances (Figure 5B). Considering that the progression category genes have highly heterogeneous expression patterns (e.g., Nieminen at al. 1998, Figure 5C), their high cell-level transcript abundance (Figure 5B) is suggestive about their critical role at the cell level. That is, progression category genes are not only highly expressed at the tissue level because they have broad expression domains, but rather because they are upregulated in individual cells irrespective of domain identity or size. These results suggest that high cell-level transcript abundance is a system-level feature of genes essential for the progression of tooth development, a pattern that seems to be shared with essential genes of single cell organisms (Dong et al., 2018). We note that although the dispensable category has several genes showing comparable expression levels with that of the progression category genes at the tissue level (Figure 2), their cell-level transcript abundances are predominantly low (Figure 5B).

Next we examined more closely the differences between progression and shape category genes, and to what extent the upregulation of the keystone genes reflects the overall expression of the corresponding pathways.

### 3.4| Keystone gene upregulation in the context of their pathways

In our data the developmental-process genes appear to have slightly elevated expression levels compared to the other protein coding genes (Figs 2, 3, 4B), suggesting an expected and general recruitment of the pathways required for organogenesis. To place the progression and shape category genes into the specific context of their corresponding pathways, we investigated in E14 mouse bulk RNAseq whether the pathways implicated in tooth development show elevated expression levels. Six pathways, Fgf, Wnt, Tgfβ, Hedgehog (Hh), Notch, and Ectodysplasin (Eda), contain the majority of progression and shape genes (Materials and Methods). First we used the RNAseq of E14 stage molars to test whether these pathways show elevated expression levels. We manually identified 272 genes belonging to the six pathways (Materials and Methods, Table S4). Comparison of the median expression levels of the six-pathway genes with the developmental-process genes shows that the pathway genes are a highly upregulated set of genes (Figure 6A, *p* < 0.0001, random resampling). This difference suggests that the experimentally identified progression and shape genes might be highly expressed partly because they belong to the developmentally upregulated pathways. To specifically test this possibility, we contrasted the expression levels of the progression and shape genes to the genes of their corresponding signaling families.

**Figure 6.**
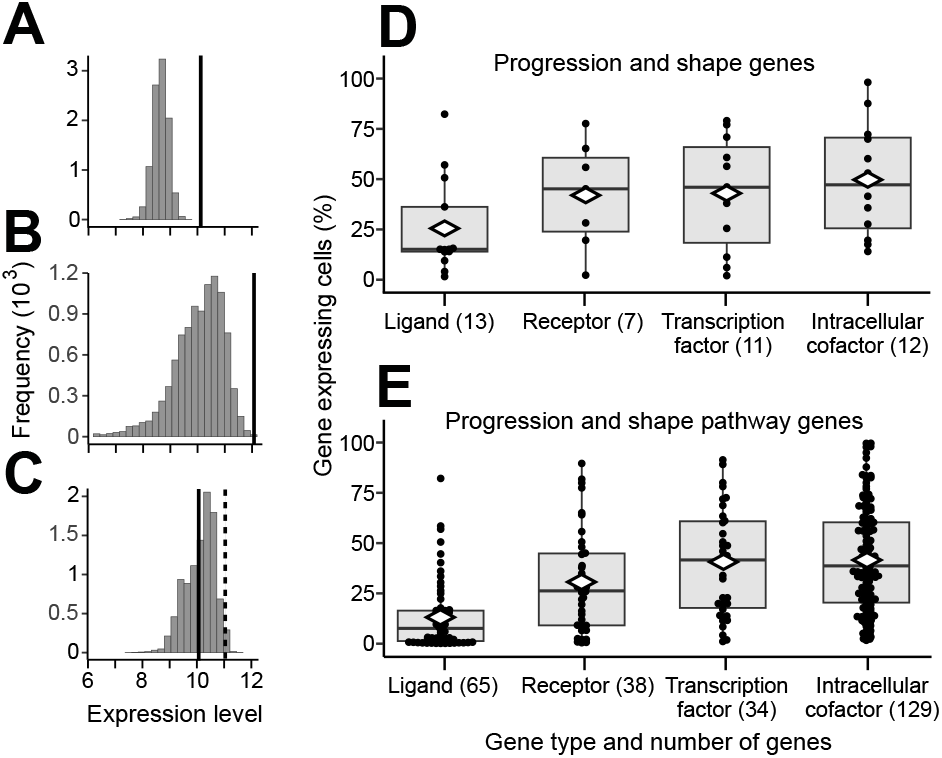
Restricted expression of ligands explains differences in upregulation. (A) Compared to all the developmental process genes, the pathways containing progression and shape category genes are generally upregulated (*p* < 0.0001, black line, random sampling using median expression levels and 10 000 permutations). (B) Median expression of progression genes (black line) shows further upregulation compared to all the genes in the corresponding pathways (*p* = 0.0004). (C) Median expression of shape category genes (black line) shows comparable expression with the genes in the corresponding pathways (*p* = 0.5919), but excluding ligands makes the shape category genes highly expressed (*p* = 0.0154, black dashed line). (D) Of the progression and shape category genes, ligands are expressed in fewer number of cells compared to the non-secreted proteins. (E) Ligands are also expressed in relatively few cells among all the pathway genes. Analyses using E14 mouse bulk RNAseq (A–C) and E14 scRNAseq (D, E). The *p* value for ligands in (D) and (E) are *p* < 0.0001 (random sampling compared to all the other gene types). Boxes in (D, E) enclose 50% of observations; the median and mean are indicated with a horizontal bar and diamond, respectively, and whiskers extend to last values within 1.5 interquartiles.

The 15 progression category genes belong to four signaling families (Wnt, Tgfβ, Fgf, Hh) with 221 genes in our tabulations. Even though these pathways are generally upregulated in the E14 tooth, the median expression level of the progression category is still further elevated (Figure 6B, *p* < 0.0001). In contrast, the analyses for the 28 shape category genes and their corresponding pathways (272 genes from Wnt, Tgfβ, Fgf, Hh, Eda, Notch) show comparable expression levels (Figure 6C, *p* = 0.5919). Whereas this contrasting pattern between progression and shape genes within their pathways may explain the subtle upregulation of the shape category (Figure 2), the difference warrants a closer look. Examination of the two gene categories reveals that compared to the progression category genes, relatively large proportion of the shape category genes are ligands (36% shape genes compared to 20% progression genes, Appendix S1). In our E14 scRNAseq data, ligands show generally smaller expression domains than other genes (roughly by half, Figure 6D, E), and the low expression of the shape category genes seems to be at least in part driven by the ligands (Figure 6C, Table S5).

Overall, the upregulation of the keystone genes within their pathways appears to be influenced by the kind of proteins they encode. In this context it is noteworthy that patterning of tooth shape requires spatial regulation of secondary enamel knots and cusps, providing a plausible explanation for the high proportion of genes encoding diffusing ligands in the shape category.

## 4| DISCUSSION

Identification and mechanistic characterization of developmentally essential or important genes have motivated a considerable research effort (e.g. Amsterdam et al., 2004; Brown et al., 2018; Bogue et al., 2018; Dickinson et al., 2016). One general realization has been that despite the large number of genes being dynamically expressed during organogenesis, only a subset appears to have discernable effects on the phenotype. This parallels with the keystone species concept used in ecological research (Paine, 1969; Terborgh, 1986). Keystone species, that may include relatively few species in a community, are thought to have disproportionally large influence on their environment. Similarly, keystone genes of development, while not necessarily essential for life, have disproportionally large effects on the phenotypic outcome of their system. Here we took advantage of this kind of in-depth knowledge on the details of the phenotypic effects of various developmental genes (Appendix S1). This allowed us to classify genes into different categories ranging from essential to ‘less essential’ and all the way to dispensable. Obviously, as in ecological data, our category groupings can be considered a work in progress as new genes and reclassifications are bound to refine the patterns. Nevertheless, our analyses should provide some robust inferences.

Most notably, genes that are essential for the progression of mouse molar development were highly expressed (Figs 2–4). These genes were highly expressed even within their pathways (Figure 6A, B) and had markedly high cell-level transcript abundances (Figure 5B). This pattern conforms to analyses of single cell organisms (Dong et al., 2018), thereby supporting expression level as one general criterion for essential genes. The high expression level of progression category genes may well signify their absolute requirement during the cap stage of tooth development. Indeed, it is typically by this stage that a developmental arrest happens when many of the progression genes are null mutated. Interestingly, mice heterozygous for the null-mutated progression genes appear to have normal teeth (Appendix S1). A possible hypothesis to be explored is to examine whether the high cell-level transcript abundance of the progression category is a form of haplosufficiency in which the developmental system is buffered against mutations affecting one allele. Another possibility, that has some experimental support (Benazet et al., 2009), is that there are regulatory feedbacks to boost gene expression to compensate for a null-allele. In contrast to the progression category, gene pairs arresting tooth development as double mutants have relatively low expression levels, perhaps suggestive that several genes in the dispensable category could be redundant to each other. This redundancy could contribute to the overall robustness of tooth development.

Because all the studied progression and shape category genes are involved in the development of multiple organ systems, our results may evolutionarily point to cis-regulatory differences that specifically promote the expression of these genes in an organ specific manner. Consequently, species that are less reliant on teeth (e.g., some seals) or have rudimentary teeth (e.g., baleen whales) can be predicted to have lowered expression levels of the progression genes. Nevertheless, at an organism level, our gene categories should not be considered as indicative of having simple effects on individual fitness. For example, producing offspring, which have defective enamel, could be more costly to the mammalian parent than offspring with a null mutation causing an arrest of tooth development with comparable defects in other organ systems, and thus early lethality. That our results may apply to other species than the mouse is supported by the similarity of the organ level expression patterns in the mouse and the rat (Figure 3), as also by the generally comparable phenotypic effects of mutations in the mouse and in the human (Nieminen, 2009). Therefore, we suggest that even though gene expression profiles may differ in details among species, the overall, high-level patterns of essential gene expression dynamics should be evolutionarily conserved.

Towards the general attempt to understand the numerous genes expressed in a developing organ system, our results point to the potential to use cell-level expression levels to identify genes critical for organogenesis. Here the single-cell transcriptomes provided a more nuanced view into the spatial patterns of the different gene categories than the tissue level transcriptomes alone, which mostly reflect the size of the expression domain (Figure 5A). In our tabulation, over a third of the shape category genes were ligands. Tooth shape patterning involves spatial placement of signaling centers that in turn direct the growth and folding of the tissue (Jernvall & Thesleff, 2012). The involvement of several secreted ligands in this patterning process, and consequently in the shape category, is likely to reflect the requirement of the developmental machinery to produce functional cusp patterns. These cusp patterns are also a major target of natural selection because evolutionary diversity of mammalian teeth largely consists of different cusp configurations. At the same time, partly due to ligands having generally more restricted expression domains compared to receptors and intracellular proteins, the shape category expression levels were found to be generally lower than that of the progression category. That ligands tend to have smaller expression domains whereas receptors have broader expression domains for tissue competence has been recognized in many unrelated studies (e.g. Bachler & Neubüser, 2001; Wessells, Grumbling, Donaldson, Wang, & Simcox, 1999; and partly in Salvador-Martínez & Salazar-Ciudad, 2015), but our analyses suggest that this is a general principle detectable in system-level transcriptome data. This pattern is also compatible with the classic concepts of tissue competence and evocators or signals produced by organizers (Waddington, 1940). Nevertheless, it remains to be explored spatially how the low signal-competence ratio emerges from highly heterogeneous expression domains of various genes, and within the complex three-dimensional context of a developing tooth (e.g., Harjunmaa et al., 2014; Pantalacci et al., 2015; Krivanek et al., 2020). In our data (for accession number, see Materials and Methods), at least the ligand *Eda* is expressed in a larger number of cells than its receptor *Edar*, suggesting that there are individual exceptions to the general pattern. Another potentially interesting observation is that *Sostdc1* and *Fst*, both secreted sequesters or inhibitors of signaling, were among the most broadly expressed of the ligands. Thus, at least some of the exceptions to the low signal-competence ratio may be modulators of tissue competence.

In conclusions, the over 20 000 genes of mammalian genomes, and even higher numbers in many plant genomes, call for systems to categorize them. Especially the high-throughput experiments have accentuated the need for comprehension of the bigger picture in genome-wide analysis. However, there is no single way to do the classification of genes. Biological complexity offers a multitude of ways to categorize, ranging from structural to functional characteristics, and from evolutionary relationships to location of expression. Here our aim was to create a categorization that would provide insight to systems-level understanding of organogenesis and still include organ level details. By combining the experimental evidence on the effects of gene null-mutations with single-cell level transcriptome data, we uncovered potential generalities affecting expression levels of genes in a developing system. With advances in the analyses of transcriptomes and gene regulation, it will be possible to explore experimental data from other organs and species to test and identify system level principles of organogenesis.

## ACKNOWLEDGEMENTS

We thank I. Salazar-Ciudad and S. F. Gilbert for advice and the members of the Center of Excellence in Experimental and Computational Developmental Biology Research for discussions. We thank A. Viherä for technical assistance. We thank P. Auvinen, L. Paulin and P. Laamanen at DNA Sequencing and Genomics Laboratory for bulk RNA sequencing. We thank J. Lahtela at FIMM Single Cell Analytics for single cell RNA sequencing.

